# Early Life Trauma Predicts Affective Phenomenology and the Effects are Partly Mediated by Staging Coupled with Lowered Lipid-Associated Antioxidant Defences

**DOI:** 10.1101/397711

**Authors:** Michael Maes, Ana Congio, Juliana Brum Moraes, Kamila Landucci Bonifacio, Decio Sabbatini Barbosa, Heber Odebrecht Vargas, Gerwyn Morris, Basant K. Puri, Ana Paula Michelin, Sandra Odebrecht Vargas Nunes

## Abstract

**Background:** Early life trauma (ELT) may drive mood disorder phenomenology, neuro-oxidative and neuro-immune pathways and impairments in semantic memory. Nevertheless, there are no data regarding the impact of ELT on affective phenomenology and whether these pathways are mediated by staging or lowered lipid-associated antioxidant defences.

**Methods:** This study examined healthy controls (n=54) and patients with affective disorders including major depression, bipolar disorder and anxiety disorders (n=118). ELT was assessed using the Child Trauma Questionnaire. In addition, we measured affective phenomenology and assayed advanced oxidation protein products; malondialdehyde, paraoxonase 1 (CMPAase) activity, high-sensitivity C-reactive protein (hsCRP), and high-density lipoprotein (HDL) cholesterol.

**Results:** ELT was associated with increased risk for mood and comorbid anxiety disorders and a more severe phenomenology, including staging characteristics (number of mood episodes), severity of depression and anxiety, suicide attempts, suicidal ideation, type of treatments received, disabilities, body mass index, smoking behaviour and hsCRP, as well as lowered health-related quality of life, socio-economic status, antioxidant defences and semantic memory. The number of mood episodes and CMPAase/HDL-cholesterol levels could be reliably combined into a new vulnerability staging-biomarker index, which mediates in part the effects of ELT on affective phenomenology, while lowered antioxidant defences are associated with increased oxidative stress. Moreover, the effects of female sex on mood disorders and affective phenomenology are mediated by ELT.

**Discussion:** The cumulative effects of different types of ELT drive many aspects of affective phenomenology either directly or indirectly through effects of staging and/or lipid–associated antioxidant defences. The results show that children, especially girls, with ELT are at great risk to develop mood disorders and more severe phenotypes of affective disorders.

## Introduction

Both major depressive disorder (MDD) and bipolar disorder (BD) are highly recurrent disorders that are characterized by increased risk for suicidal behaviours [1-3], lowered health-related quality of life (HR-QoL), increased disabilities [4-7], lowered socio-economic status [8-11], smoking behaviours [12-14], increased incidence of metabolic syndrome [15] and cognitive impairments including semantic memory deficits [16].

There is also evidence that neuro-immune, neuro-oxidative and neuro-nitrosative pathways play a key role in the pathophysiology of both MDD and BD [17-20]. Both mood disorders are associated with lowered lipid-associated antioxidant defences, which protect against the detrimental effects of reactive oxygen species (ROS), including lowered lecithin cholesterol acyltransferase (LCAT) and paraoxonase (PON1) activities and lowered high-density lipoprotein (HDL)-cholesterol, vitamin E and coenzyme Q^10^ levels [21-30]. These lowered lipid-associated antioxidant defences contribute to increased ROS production and consequent damage to lipid membranes, including omega-3 long-chain polyunsaturated fatty acids [30-33] and proteins [34,35]. Such reactions may lead to the production of advanced oxidation protein products (AOPP) [34,35], generation of neoepitopes including malondialdehyde (MDA) and azelaic acid, and IgM-mediated autoimmune responses directed against those neoepitopes [20,30,36-41]. Lowered PON1 (CMPAase) activity is tightly coupled with lowered HDL-cholesterol levels and number of depressive and (hypo)manic episodes [29]. Activated immune-inflammatory pathways, which are tightly connected with enhanced oxidative stress pathways, are other hallmarks of mood disorders [20], and include increased levels of pro-inflammatory cytokines and acute phase proteins, such as C-reactive protein (CRP) [17,42]. Nevertheless, the sources of these activated oxidative and immune pathways are not completely elucidated, although leaky gut, leaky teeth and psychological stressors are involved [34,43,44].

Early life trauma (ELT), including physical and emotional neglect or abuse and sexual abuse as well as parental discord and bullying, is an environmental factor associated with the onset of affective disorders [45-52]. ELTs impact the course of illness and may cause more recurrent episodes [53,54], increase severity of illness and suicidal behaviours [51,55,56], increase comorbid anxiety disorders [57,58] and obesity [59] and delayed neurocognitive development [60]. Women show a higher incidence of affective disorders and higher scores on severity of illness as compared with men [61-63], whilst women also report more ELTs than men [64]. Nevertheless, it has remained elusive whether the effects of female sex on affective phenomenology are mediated by ELTs, and whether the effects on affective phenomenology are mediated by staging characteristics.

There is also evidence that ELTs may sensitize and activate neuro-immune and neuro-oxidative pathways. For example, in mood disorders, physical neglect was significantly associated with indicants of lipid peroxidation, nitro-oxidative stress and lipid and protein oxidation, while sexual abuse was accompanied by lowered antioxidant levels including zinc, albumin and SH groups [65]. ELTs are positively associated with increased pro-inflammatory cytokines and CRP, while these aberrations may persist in euthymic phases of BD [review: 66]. In another study, increased levels of hsCRP were significantly associated with sexual abuse rather than with BD, whilst the differences in hsCRP between BD patients and controls disappeared after controlling for body mass index (BMI) [66].

Hence, the aim of this study was to examine whether ELTs are associated with mood disorders, comorbid anxiety, staging characteristics (number of mood episodes and lifetime suicidal attempts), current suicidal ideation, severity of illness, type of treatments, increased disabilities and lowered HR-QoL, socio-economic variables (income and education), BMI, smoking behaviour, semantic impairments, lipid-associated antioxidant defences (HDL-cholesterol and PON1 activity), lipid/protein oxidation and hsCRP. Secondly we examine whether the effects of female sex on affective phenomenology are mediated by ELTs and whether the effects of ELTs on affective phenomenology are mediated by staging characteristics or biomarkers.

## Methods

### Study population

This study examined ELT and first-degree relative histories of mental disorders in 54 controls and 118 participants with affective disorders recruited at the outpatient Psychiatric Clinics at the University Hospital of the Universidade Estadual de Londrina (UEL), Parana, Brazil. We included participants aged 18 to 65 years, either sex, and from all ethnicities (self-declared). The controls were recruited by word of mouth from the same catchment area as the patients, namely Parana, Brazil. We included patients with MDD (n=37), BD (n=68) and DSM-IV-TR anxiety disorders (n=81, namely generalized anxiety disorder (GAD), social phobia, simple phobia and panic disorder). Patients with mood disorders were in (partial) remission and the index episode was not of (hypo)manic polarity. All participants provided written informed consent to take part in the study, with the experimental procedures being approved by the Research Ethics Committee at UEL (protocol number: CAAE 34935814.2.0000.5231).

We excluded the following participants: pregnant women, participants with mental disorders other than BD, MDD or anxiety disorders, including schizophrenia, psycho-organic disorders, neuro-inflammatory and neurodegenerative disorders, including multiple sclerosis, Parkinson and Alzheimer’s disease, participants who were taking omega-3 polyunsaturated fatty acids or antioxidant agents during the past four weeks prior to inclusion in the study, and subjects who had ever taken non-steroidal anti-inflammatory drugs, interferon or glucocorticoids. Furthermore, we excluded subjects with medical conditions that are characterized by immune activation or immunosuppression, such as hepatitis B and C virus infection, HIV infection, acute and chronic renal failure, inflammatory bowel disease, chronic obstructive pulmonary disease, rheumatoid arthritis and type I diabetes.

### Clinical assessments

Patients and controls completed a semi-structured interview, which comprised socio-demographic data, education (in years), employment status, income (in reais/month) and marital status. The interview also included (a) the number of depressive and manic episodes, and total number of episodes; and (b) the use of medication, including antidepressants, antipsychotics, lithium and anticonvulsant stabilizers.

ELT was assessed using the Childhood Trauma Questionnaire (CTQ). This is a self-rating scale for adolescents and adults investigating a history of physical and emotional abuse or neglect and sexual abuse during childhood; the frequency of each ELT is rated on a five-point Likert scale applied to 28 items [67,68]. We used a Portuguese version of the CTQ instrument validated for use in the Brazilian population [68].

Diagnoses of MDD, BD, and anxiety disorders were made by trained research psychiatrists using the Structured Clinical Interview for the DSM-IV-TR (clinical version) translated and validated for application in Brazilian samples [69]. Severity of depression was assessed employing the 17-item Hamilton Depression Rating Scale (HAMD) [70], translated and validated for the Brazilian population [71]. Severity of anxiety was assessed employing the Hamilton Anxiety Rating Scale (HAMA) [72]. We used the Columbia–Suicide Severity Rating Scale (C-SSRS) to rate severity of current suicidal ideation and number of prior suicide attempts [73]. The Clinical Global Impressions (CGI) scale was used to assess overall severity of illness, which scores the subject status on a scale from 1 to 7 [74]. We used the Sheehan Disability Scale to assess disabilities. This self-rating scale scores occupational, social life, leisure, family life, activities, and household activities [75]. The WHO Quality of Life Instrument-Abbreviated Version (WHO-QOL-BREF) [76] was used in a validated Brazilian Portuguese translation [77]. We used the sum of the four WHO-QoL domains, namely physical health, psychological health, social relationships and environment. A subset of the participants (n=83) also completed the Verbal Fluency Test (VFT), animal category. We used DSM-IV-TR criteria to make the diagnosis of tobacco use disorder (TUD). Body mass index (BMI) was calculated as body mass (kg) divided by square of height (m^2^).

### Biomarker measurements

Peripheral blood was sampled at 8 a.m. after 12 h fasting, on the same day as the semi-structured interview and clinical ratings. We assessed the levels of hsCRP, MDA, AOPP, PON1 activity, and HDL-cholesterol. MDA levels were assessed using both the thiobarbituric acid (TBA) assay, in which MDA reacts with TBA to generate the colored (TBA)_2_-MDA adduct, and by high-performance liquid chromatography (HPLC; Alliance e2695, Waters’, Barueri, SP, Brasil) [78]. Experimental conditions included the use of a column Eclipse XDB-C18 (Agilent, USA); mobile phase consisting of 65% phosphate buffer (50 nM pH 7.0) and 35% HPLC-grade methanol; flow rate of 1.0 mL/minute; temperature of 30 °C; wavelength of 532 nm (the absorption frequency of (TBA)_2_-MDA). MDA concentration in the samples was quantified based on a calibration curve and was expressed in μmol MDA/mg proteins. AOPP was quantified spectrophotometrically using the method described by Hanasand et al. [79] using a microplate reader, model EnSpire (Perkin Elmer, Waltham, MA, EUA) with an absorbance wavelength of 340 nm. AOPP concentration was expressed in chloramine units (μM). PON1 activity was determined using 4-(chloromethyl)phenyl acetate (CMPA; Sigma, USA), which is an alternative to the use of the toxic paraoxon [29]. The analysis was conducted in a spectrophotometer microplate reader (EnSpire, Perkin Elmer, USA). All assays were carried out in triplicate and replicates that varied by 10% or greater were repeated. The CMPAase activity levels were adjusted for PON1 genotypes using regression analysis. HDL-cholesterol was assayed by automated methods using the Dimension RxL (Deerfield, IL, USA). hsCRP was assayed using a nephelometric assay (Behring Nephelometer II, Dade Behring, Marburg, Germany). The inter-assay coefficients of variability were less than 10%.

### Statistics

Analysis of variance (ANOVA) was used to assess differences in scale variables (e.g. clinical, socio-demographic and biomarker data) between categories, while analysis of contingency tables (χ^2^-test) was used to check associations between nominal variables. Binary logistic regression analysis was used to check the significant explanatory variables predicting binary outcome variables, including BD (versus no BD), any anxiety disorders (versus no anxiety disorder) and TUD (with no-TUD as a reference group). All tests were two-tailed and an alpha level of 0.05 was considered to be statistically significant. All analyses were performed using the IMB-SPSS software version 24 for Windows. We used SmartPLS (Partial Least Squares) analysis, a structural equation modeling technique coupled with path modeling using PLS-structural equation modeling algorithms [80] to examine the effects of ELTs combined with first-degree relative histories (both entered as input variables) on affective phenomenology (output variables). Variables were included as single indicators (e.g. BMI, income) or as latent vectors (LV) extracted from indicator variables in reflective models (e.g. the four WHO-QoL domain scores reflecting general HR-QoL). PLS path analysis was only performed when the model and constructs complied with specific quality criteria, namely: (a) model fit standardized root mean residual (SRMR) < 0.08 [81]; (b) composite constructs have good discriminant validity and reliability as indicated by average variance extracted (AVE) > 0.5; Cronbach’s alpha > 0.7, composite reliability > 0.7 and rho_A > 0.8; (c) indicators are only included in LVs when the factor loadings are > 0.45 with p < 0.01; (d) construct cross-validated redundancies and communalities are checked [81]. Consequently, path coefficients with t-values and exact p-values, total effects, total indirect and specific indirect effects and t-values for the outer model were computed.

## Results

### Construction of an ELT LV and staging LV

In order to examine whether a first LV extracted from the five ELT data (ELT-LV) performed well as a composite score we used PLS factor analysis and calculated composite reliability data. The LV extracted from the 5 ELT data shows that ELT-LV has an adequate Cronbach alpha (0.848), composite reliability (0.893) and rho_A values (0.877), while the AVE value was very good (0.630). All 5 items scored highly on this LV (all >0.587). In order to examine whether staging and lipid-associated antioxidant defences (HDL-cholesterol and PON1/residualized CMPAase activity) could be reliably combined in one underlying construct we carried out PLS factor analysis. This showed that the LV extracted from number of depressive episodes, number of hypomanias, CMPAase and HDL-cholesterol showed a good composite reliability (0.808), Cronbach alpha (0.734), rho_A (0.860) and AVE (0.534), while all four variables loaded significantly (> 0.484) on this LV and were significant at the p < 0.01 level. This LV was named “EPIBIOL-LV” (from <u>epi</u>sodes coupled with <u>bio</u>markers).

### Characteristics of the ELT-LV subgroups

In order to show the measurements of the clinical and biological data in ELT-LV subgroups, we divided the participants into three equal-sized groups based on 33.3333 and 66.6666 percentiles of the ELT-LV scores. Table 1 shows the ELT differences between the three groups. Sexual and physical abuse were significantly increased in the high ELT-LV group as compared with the low and medium ELT-LV groups. Scores on emotional abuse and emotional and physical neglect were significantly different between the three groups and increased from the low → medium → high ELT-LV groups.

**Table 1.** Measurements of five early life trauma (ELT) in subjects divided into high, medium and low values of ELT-latent vector (LV) scores

| Variables | ELT-LV <<br>-0.6693 <sup>A</sup> | -0.6693 < ELT-LV <<br>+0.2217 <sup>B</sup> | ELT-LV<br>≥+0.2217 <sup>C</sup> | F/X<sup>2</sup> | df | p |
| --- | --- | --- | --- | --- | --- | --- |
| Sexual abuse | 5.09 (0.43) <sup>C</sup> | 5.62 (1.57) <sup>C</sup> | 8.70 (5.21) <sup>A,B</sup> | 22.02 | 2/169 | <0.001 |
| Physical abuse | 5.75 (1.06) <sup>C</sup> | 7.05 (2.37) <sup>C</sup> | 13.02 (4.32) <sup>A,B</sup> | 101.33 | 2/169 | <0.001 |
| Emotional abuse | 5.63 (0.92) <sup>B,C</sup> | 9.00 (2.76) <sup>A,C</sup> | 16.56 (4.51) <sup>A,B</sup> | 186.48 | 2/169 | <0.001 |
| Emotional neglect | 6.30 (1.65) <sup>B,C</sup> | 9.88 (3.94) <sup>A,C</sup> | 17.39 (4.18) <sup>A,B</sup> | 153.35 | 2/168 | <0.001 |
| Physical neglect | 5.63 (1.19) <sup>B,C</sup> | 7.47 (2.53) <sup>A,C</sup> | 11.40 (3.77) <sup>A,B</sup> | 67.74 | 2/169 | <0.001 |

### Effects of ELT-LV and EPIBIOL-LV on affective phenomenology

Consequently, we constructed a PLS path model as shown in **Figure 1** with affective phenomenology as outcome variables (diagnosis, disabilities, HR-QoL, CGI, severity, suicidal attempts) and ELT-LV and EPIBIOL-LV as explanatory variables, while sex was also an explanatory variable for ELT-LV and EPIBIOL-LV and the latter for ELT-LV. The five Sheehan subdomain scores were used as indicators for the “disabilities LV” (the loading of Sheehan 4 subdomain was not significant and thus deleted from the analysis), the four WHO-QoL subdomain scores were employed as indicators for the WHO-QoL LV, HAMA and HAMD as indicators for severity of illness and mood disorders versus controls, any anxiety disorder and BD as indicators for a “diagnosis LV”. Figure 1 shows the outcome of this PLS path model with path coefficients (pc) and exact p-values. The model quality data were excellent with model fit SRMR = 0.064, composite reliabilities > 0.8, Cronbach alpha > 0.7, AVE values > 0.5 and rho_A values > 0.8. We found that a large part of the variances in CGI (22.5%), WHO-QOL (50.0%), disabilities (32.2%), severity LV (32.1%), income (11.8%) and diagnosis (62.7%) was explained by effects of EPIBIOL-LV and ELT-LV combined. All these associations remained significant after considering possible intervening effects of BMI and TUD (entered as additional input variables). EPIBIOL-LV had always a greater impact on these outcome variables than the “BIOL-LV” (extracted from the two HDL-cholesterol and PON1 indicators) or “episodes LV” (number of episodes) alone. For example, the impact of EPIBIOL-LV (pc = 0.317, p < 0.001, see Figure1) on “disabilities LV” was greater than the impact of BIOL-LV (pc = −0.284, p < 0.001) or episodes LV (pc = −0.220, p = 0.007) alone. Lifetime suicidal attempts (24.7%) and current suicidal ideation (12.7%) were associated with EPIBIOL-LV, although number of episodes, and not antioxidant levels, were the significant predictors. There was a significant effect of ELT-LV, but not EPIBIOL-LV, on BMI. There were specific indirect effects of ELT-LV on CGI (t = 2.99, p = 0.003), disabilities (t = 3.36, p = 0.001), WHO-Qol (t = −4.62, p < 0.001), severity (t = 3.32, p = 0.001), suicidal ideation (t = 3.51, p < 0.001), suicide attempts (t = 4.40, p < 0.001), income (t = −2.12, p = 0.034) and diagnosis (t = 4.32, p < 0.001). There were highly significant total effects of ELT-LV on EPIBIOL-LV (t = 5.70, p < 0.001) and on all variables shown in Figure 1.

The BIOL-LV was significantly associated with income (pc = 0.179, p = 0.017), severity LV (pc = −0.288, p < 0.001), CGI (pc = −0.307, p < 0.001), disabilities LV (pc = −0.284, p < 0.001), WHO-QoL LV (pc = 0.277, p < 0.001), diagnosis LV (pc = −0.236, p < 0.001) and an “oxidation LV” with AOPP and MDA as indicator variables (pc = −0.235, p < 0.001). There were no associations between this BIOL-LV and BMI, suicidal ideation, suicidal attempts, and use of lithium, antidepressants, mood stabilizers, and antipsychotics. All those effects remained significant after controlling for effects of TUD.

Binary logistic regression analysis showed that ELT-LV was significantly associated with mood disorders (W = 22.21, df = 1, p < 0.001, OR = 3.15, 95%CI: 1.96-5.09); BD (W = 21.41, df = 1, p < 0.001, OR = 2.35, 95%CI: 1.64-3.37); any anxiety disorder (W = 15.65, df = 1, p < 0.001, OR = 2.03, 95%CI: 1.43-2.88); employment (W = 9.81, df = 1, p = 0.002, OR = 0.60, 95%CI: 0.43-0.82); use of mood stabilizers (W = 15.48, df = 1, p < 0.001, OR = 2.14, 95%CI: 1.47-3.13); use of antipsychotics (W = 9.35, df = 1, p = 0.002, OR = 1.77, 95%CI: 1.23-2.54) and lithium (W = 11.83, df = 1, p = 0.001, OR = 2.00, 95%CI: 1.35-2.96), but not TUD and use of antidepressants.

### Effects of sex on affective phenomenology are mediated by ELT-LV

Table 2 shows that there is a significant association between sex and the ELT-LV groups, with significantly more women in the high ELT-LV group. Therefore, we examined the specific indirect effects of sex on affective phenomenology mediated by ELT-LV and sex was added as a direct input variable for ELT-LV and EPIBIOL-LV in the PLS analysis shown in Figure 1. There was a significant direct effects of sex on ELT-LV, but not on the EPIBIOL-LV. We found significant total effects of sex (male = 1, female = 0) on ELT-LV (t = −3.45, p = 0.001), EPIBIOL-LV (t = −2.76, p = 0.006), severity of illness (t = −2.61, p = 0.009), suicidal ideation (t = −2.26, p = 0.024), suicide attempts (t = −2.52, p = 0.012), CGI (t = −2.80, p = 0.005), disabilities (t = −2.84, p = 0.005), WHO-QoL score (t = 3.00, p = 0.003), income (t = 2.18, p = 0.030), diagnosis LV (t = −3.20, p = 0.001) and BMI (t = −1.98, p = 0.048). All these sex effects were mediated by ELT-LV and/or the path from ELT-LV to EPIBIOL-LV.

**Table 2.** Socio-demographic and clinical data of subjects divided into high, medium and low values of an early life trauma (ELT)-latent vector (LV)

| Variables | ELT-LV < -0.6693 <sup>A</sup> | -0.6693 < ELT-LV < +0.2217 <sup>B</sup> | ELT-LV ≥ +0.2217 <sup>C</sup> | F/X<sup>2</sup> | df | p |
| --- | --- | --- | --- | --- | --- | --- |
| Age (years) | 41.8 (11.2) | 44.4 (11.8) | 42.2 (10.3) | 0.89 | 2/169 | 0.414 |
| Sex (female / male) | 37 / 20 <sup>C</sup> | 41 / 17 <sup>C</sup> | 51 / 6 <sup>A,B</sup> | 10.04 | 2 | 0.007 |
| Education (years) | 12.4 (6.1) <sup>C</sup> | 11.3 (5.0) | 9.8 (4.1) <sup>A</sup> | 3.56 | 2/168 | 0.031 |
| Income (reais/month) | 3.8 (1.5) <sup>C</sup> | 3.3 (1.4) | 2.9 (1.4) <sup>A</sup> | 4.92 | 2/167 | 0.008 |
| Employed (No/yes) | 14 / 42 <sup>B,C</sup> | 27 / 31 <sup>A</sup> | 32 / 25 <sup>A</sup> | 11.73 | 2 | 0.003 |
| Single/Married/separated-widowed | 18 / 29 / 10 | 13 / 34 / 11 | 7 / 34 / 16 | 7.01 | 4 | 0.136 |
| Number depressions | 1.48 (3.98) <sup>C</sup> | 2.63 (3.35) <sup>C</sup> | 5.64 (4.79) <sup>A,B</sup> | 15.06 | 2/160 | <0.001 |
| Number (hypo)mania | 0.47 (1.64) <sup>B,C</sup> | 3.43 (5.75) <sup>A</sup> | 4.12 (6.45) <sup>A</sup> | 8.30 | 2/169 | <0.001 |
| HAMD | 4.0 (5.8) <sup>B,C</sup> | 6.7 (7.9) <sup>A,C</sup> | 10.7 (6.3) <sup>A,B</sup> | 18.00 | 2/169 | <0.001 |
| HAMA | 7.0 (7.9) <sup>B,C</sup> | 11.5 (9.8) <sup>A</sup> | 17.1 (9.7) <sup>A</sup> | 16.15 | 2/158 | <0.001 |
| Number suicide attempts | 0.65 (3.15) <sup>C</sup> | 0.52 (1.35) <sup>C</sup> | 2.18 (3.74) <sup>A,B</sup> | 5.68 | 2/169 | 0.004 |
| Current suicidal ideation (No/Yes) | 50 / 7 <sup>C</sup> | 56 / 2 <sup>C</sup> | 36 / 21 <sup>A,B</sup> | 23.84 | 2 | <0.001 |
| No mood disorder / BD / MDD | 39 / 7 / 11 <sup>B,C</sup> | 19 / 26 / 13 <sup>A</sup> | 9 / 35 / 13 <sup>A</sup> | 39.34 | 4 | <0.001 |
| No anxiety / Anxiety disorders | 42 / 15 <sup>C</sup> | 31 / 27 | 18 / 39 <sup>A</sup> | 20.29 | 2 | <0.001 |
| CGI | 2.36 (1.63) <sup>B,C</sup> | 3.30 (1.36) <sup>A,C</sup> | 3.95 (1.17) <sup>A,B</sup> | 18.41 | 2/166 | <0.001 |
| Total Sheehan score | 5.3 (8.4) <sup>B,C</sup> | 11.2 (9.5) <sup>A,C</sup> | 15.7 (9.8) <sup>A,B</sup> | 18.02 | 2/168 | <0.001 |
| Total Who-QoL score | 93.1 (13.1) <sup>B,C</sup> | 84.8 (15.0) <sup>A,C</sup> | 71.4 (14.7) <sup>A,B</sup> | 33.51 | 2/168 | <0.001 |
| No TUD / TUD | 31 / 26 | 25 / 33 | 21 / 36 | 3.65 | 2 | 0.162 |
| Body mass index (kg/m<sup>2</sup>) | 25.7 (4.9) | 26.2 (4.4) | 27.6 (5.2) | 2.31 | 1/161 | 0.103 |

All results are shown as mean ( ± SD)
<sup>A,B,C</sup>: denote pair-wise differences between the three groups
HAMD: Hamilton Depression Rating Scale; HAMA: Hamilton Anxiety Rating Scale; BD: bipolar disorder; MDD: major depressive disorder; CGI: clinical global impression; WHO-QoL: WHO-Quality of Life; TUD: tobacco use disorder.

**Table 3.** Biomarker data and verbal fluency test (VFT) results in subjects divided into high, medium and low values of an early life trauma (ELT) latent vector (LV)

| Variables | ELTFA-HOC <<br>-0.6693 <sup>A</sup> | -0.6693 < ELTFA-HOC < +0.2217 <sup>B</sup> | ELTFA-HOC<br>≥+0.2217 <sup>C</sup> | F | df | p |
| --- | --- | --- | --- | --- | --- | --- |
| HDL-cholesterol (mg/dL) | 51.0 (17.0) | 48.0 (15.7) | 46.2 (12.8) | 1.32 | 2/156 | 0.270 |
| Residualized CMPAase (z scores) | 0.25 (0.96) | -0.12 (1.03) | -0.13 (0.98) | 2.40 | 2/147 | 0.095 |
| MDA (mmol/mg of protein)* | 63.0 (20.1) | 67.2 (24.2) | 64.0 (21.7) | 0.42 | 2/146 | 0.656 |
| AOPP (μM)* | 73.2 (46.2) | 85.9 (50.5) | 83.9 (45.2) | 1.05 | 2/147 | 0.353 |
| hsCRP (mg/dL)* | 3.0 (3.0) | 4.8 (7.5) | 4.2 (4.5) | 1.11 | 2/157 | 0.332 |
| EPIBIOL-LV | -0.48 (0.81) <sup>B,C</sup> | +0.01 (0.83) <sup>A,C</sup> | +0.49 (1.12) <sup>A,B</sup> | 15.46 | 2/168 | <0.001 |
| VFT words / minute | 16.5 (5.3) <sup>C</sup> | 13.2 (3.6) | 13.1 (3.6) <sup>A</sup> | 4.06 | 2/80 | 0.021 |

### Effects of ELT-LV on biomarkers

We examined whether hsCRP, AOPP or MDA could be combined with the staging data in a reliable LV (as with PON1 activity and HDL-cholesterol in Table 1). These three biomarkers could not be combined with the “episodes LV” to make reliable composite scores. We also examined whether these three biomarkers had comparable effects as EPIBIOL-LV on the clinical outcome data and detected that only AOPP had some effects, namely on CGI (t = 4.53, p < 0.001), WHO-QoL (t = −2.56, p = 0.011) and severity of illness (t = 2.55, p = 0.011).

In order to examine the effects of ELT-LV on the biomarkers we performed a further PLS path analysis as shown in **Figure 2**, with HAMD, total Sheehan and total HR-QoL scores as indicators of an “outcome LV” and AOPP/MDA LV, HDL/CMPAase LV, episodes LV, hsCRP and ELT-LV as explanatory variables, while adjusting for TUD, BMI and sex, whereby sex was also an explanatory variable for ELT-LV. The model quality data were excellent with model fit SRMR = 0.045, composite reliability values > 0.8, Cronbach alpha values > 0.8, AVE > 0.5 and rho_A values > 0.8. We found that 54.7% of the variance in the outcome LV was explained by significant effects of AOPP/MDA LV, episodes LV, ELT-LV, TUD and female sex. 22.0% of the variance in AOPP/MDA LV was explained by HDL/CMPAase LV, sex and BMI, while 17.6% of the variance in HDL/CMPAase LV was explained by episodes LV, TUD and BMI. 27.2% of the variance in hsCRP was explained by BMI and sex. There were total effects of ELT-LV on hsCRP (t = 2.51, p = 0.013), AOPP/MDA LV (t = 2.27, p = 0.024) and HDL/CMPAase LV (t = −3.39, p = 0.001). There were total indirect effects of ELT-LV on AOPP/MDA (t = 2.27, p = 0.024). The effects of ELT-LV on HDL/CMPAase was mediated by BMI (t = −2.00, p = 0.015) and episodes LV (t = −2.43, p = 0.015), while the effects of ELT-LV on CRP were mediated by BMI (t = 2.51, p = 0.013).

### Effects of ELT-LV on VFT

**Figure 3** examines the effects of ELT-LV on VFT considering that severity of illness and staging could affect VFT. There were no significant effects of any of the biomarkers on VFT and therefore these indicators were eliminated from the model. Figure 3 shows a PLS model with VFT as final outcome and severity LV, episodes LV, education and ELT-LV as input variables while adjusting for age and sex. The model quality data were again excellent with SRMR = 0.058, and Cronbach alpha values > 0.843, rho_A > 0.881, composite reliabilities > 0.856 and AVE > 0.553. We found that 20.0% of the variance in VFT was explained by episodes LV, education and sex, while severity LV was not significant. There were specific indirect effects of ELT-LV on VFT, which were mediated by episodes LV (t = −2.55, p = 0.010).

## Discussion

### ELT strongly predicts affective phenomenology

The first major finding of this study is that increased ELT-LV scores are highly significantly associated with mood disorders, in particular with BD, and many characteristics of affective phenomenology, including severity of depression and anxiety, staging characteristics including number of mood episodes, suicidal behaviours including number of lifetime suicide attempts and current suicidal ideation, the presence of comorbid anxiety disorders and type of treatments received namely use of mood stabilizers, lithium and antipsychotics. Cross-sectional and longitudinal studies show that ELT and especially multiple ELTs or the accumulation of different traumatic events in childhood are associated with the development of depression later in life [82-84]. A recent review shows that childhood ELTs and especially multiple ELTs also increase risk for developing BD and more severe phenotypes of BD [64,85-87]. Our findings also extent those of previous papers showing that ELT may be associated with increased severity of illness and more suicidal behaviours [55,56], recurrent episodes [88], and comorbid anxiety disorders [57,58,89]. In addition, we found that ELT-LV has a strong impact on lowered income, increased disabilities and a lowered HR-QoL. A prospective study showed that there were only weak connections between ELT and lowered QoL in later life [90], while in BD patients, ELT is accompanied by lowered HR-QoL [91]. Moreover, we found that ELT-LV also affected BMI, but not smoking behaviour. Previously, it was reported that among individuals with psychopathology, ELT may predispose to adult cigarette smoking [92]. Previous research also showed a connection between ELT and increased metabolic burden. For example, in first-episode psychosis patients, those with ELT had higher BMI when compared with healthy controls [93], while ELT is also associated with obesity [94].

### The effects of ELT on affective phenomenology are mediated by staging and lowered antioxidant defences

The second major finding of this study is that the effects of the vulnerability factor ELT-LV on affective phenomenology are in part mediated by recurrent episodes and lipid-associated antioxidant defences, namely lowered HDL-cholesterol and CMPAase activity. In addition, there was a significant association between recurrent episodes and lowered levels of these lipid-associated antioxidants so that a latent vector could be computed reflecting the combined effects of episode recurrence (EPI) and these biomarkers (BIOL), named EPIBIOL-LV. This LV reflects the shared variance between increased staging and lowered antioxidant defences in affective disorders. While both type of variables have significant effects on affective phenomenology their combination showed stronger effects.

Previously, we have reported that there is a strong association between recurrence of depressive and (hypo)manic episodes and lowered CMPAase activity, indicating that sensitization mechanisms are involved in lowered levels of these antioxidants [29]. Given the cross-sectional nature of our study, it is difficult to conclude whether lowered antioxidant defences are a primary phenomenon or occur as a consequence of staging. Nevertheless, patients with drug-naïve first-episode psychosis (of whom part develop BD) show lowered PON1 activity levels, indicating that lowered PON1 activity is already present during a first episode [95]. Thus, the most parsimonious explanation is that staging, lowered antioxidant defences and shared variance among staging and lowered antioxidant defences contribute to affective phenomenology.

We have discussed previously that lowered levels of PON1 or CMPAase activity and lowered HDL-cholesterol levels are tightly coupled phenomena in mood disorders [96]. Thus, PON1 is integrated into HDL molecules and protects HDL, low-density lipoprotein and lipids against oxidation. Mood disorders are accompanied by immune-inflammatory responses including activated macrophages that produce nitric oxide (NO) and myeloperoxidase (MPO), thereby generating HCl and peroxynitrite, which in turn may attenuate PON1 binding to HDL and damage and inactivate PON1 in HDL [97]. This downregulation of the PON1/CMPAase - HDL functional complex is accompanied by increased MPO activity and thus increased production of HCl and peroxynitrite which may generate more lipid and protein oxidation as measured with MDA and AOPP [96,97].

Previously, we have reported that in mood disorders and first episode psychosis, lowered PON1-associated antioxidant defences are associated with signs of increased oxidation and immune activation as indicated by increased levels of interleukin (IL)-6, IL-4 and IL-10 [19,29,95,96]. Here, we found that lowered HDL-PON1 antioxidant defences are associated with increased lipid and protein oxidation and consequently with severity of illness and that increased AOPP levels additionally predict CGI and lowered HR-QoL. Interestingly, in the current study we found that the significant effects of ELT-LV on hsCRP levels were completely mediated by BMI. These data extend those of previous reports that plasma hsCRP is strongly determined by BMI [66]. Most importantly, while lowered antioxidant defences and increased oxidative stress biomarkers affected the outcome of affective disorders, hsCRP did not have any effect.

All in all, this study established two new paths: (a) from the vulnerability factor ELT-LV → a combined staging-biomarker → increased lipid/protein oxidation → further aggravation of the affective phenotype; and (b) from the vulnerability factor ELT-LV → increased BMI → increased hsCRP. Therefore, we may conclude that while lipid-associated antioxidant defences and lipid/protein oxidation play a role in the outcome of affective disorders, hsCRP is not involved, although it may be increased as a consequence of increased BMI that may occur in affective disorders.

### Effects of female sex on affective phenomenology

The third major finding of this study is that female sex was associated with an increased incidence of affective disorders, affective phenomenology and with higher ELT-LV scores. Firstly, we found that female sex increased severity of illness, suicidal ideation, suicide attempts, disabilities, use of psychotropic medications, and BMI and lowered WHO-QoL score and income. It is well known that females are twice as likely as males to develop mood and comorbid anxiety disorders and have higher scores on severity of illness and more affective symptoms [61-63]. These sex-differences are established across cultures even after controlling for help-seeking behaviors, suggesting biological differences [98,99].

Secondly, we found that the significant effects of sex on affective phenomenology are mediated by ELT-LV, suggesting that sex may drive affective phenomenology by modulating ELT. The results indicate that girls are more likely to suffer from childhood ELT and that this sex-interaction may explain the effects of sex on affective phenomenology. These results extend those of Etain et al. [100] who reported that affective phenomenology is modulated by a sex non-specific effect of ELT in that women with BD report more ELT than men, and women show stronger associations between ELT and BD characteristics, including number of depressive episodes and suicide attempts [100]. In this respect, it is interesting to note that girls being victimized by bullies at six years old are more likely than boys to remain a victim four years later [101]. Girls report more childhood physical abuse than boys, while ELT may have a more profound impact on girls than boys [102]. The latter authors reported that such effects may be related to sex differences in the insula’s anterior circular sulcus. In our study, however, female sex appears to protect against lowered lipid-associated antioxidant defences and increased oxidation and therefore these neuro-oxidative pathways do not explain the effects of female sex on affective phenomenology.

### ELT-LV and deficits in semantic memory

The fourth major finding of this study is that ELT-LV also predicts impairments in verbal fluency and that this effect is mediated by staging of illness. There is a vast literature that mood disorders are characterized by cognitive impairments including semantic memory [16,103]. There is now evidence that ELT predisposes towards cognitive dysfunctions including in the domains of memory, executive functioning and emotion regulation [60,104].

## Conclusions

A combination of five different ELTs appear to drive many aspects of affective phenomenology especially in women, indicating that children, especially girls, who experience ELT are at an increased risk to develop affective disorders with a more severe psychopathology. Moreover, the effects of ELT on affective phenomenology are in part mediated by staging and/or lowered lipid–associated antioxidant defences. The newly constructed latent vector, which reflects shared variance among staging and PON1 and HDL-associated antioxidant defences, is a second vulnerability factor that together with ELT is associated with affective phenomenology. The findings show that staging-associated antioxidant defences play an important role in affective disorders. Moreover, other lipid-associated antioxidants systems may play a role in affective disorders, including lowered levels of LCAT, vitamin E, coenzyme Q_10_, glutathione and glutathione peroxidase. Therefore, ELT should be treated in the earliest phase of illness, whilst lipid-associated antioxidant defences are another target which could prevent further relapses. Recently, we have reviewed different strategies which may enhance PON1 activity, including administration of polyphenols, vitamin E/C and oleic acid, while also a Mediterranean diet and physical exercise and statins and fibrates may enhance PON1 activity [105]. Future research should (a) discover new treatments targeting PON1 binding to HDL to improve protection against activated neuro-oxidative and neuro-immune pathways and to remove consequent oxidative injuries from the body; and (b) examine the association between staging and antioxidant defences in prospective studies as well as the effects of ELT on these factors.

## Acknowledgements

The authors wish to thank the Centre of Approach and Treatment for Smokers, Psychiatric Unit at UEL, Clinical Laboratory of the University Hospital and Laboratory of Research and Graduate College Hospital (LPG), Brazil.

## Authorships

All authors contributed to the writing up of the paper. The work was designed by SOVN, MM, DSB, JBM and HOV. Data were collected by SOVN, HOV, AC, and JBM. Laboratory analyses were conducted by KLB, APM and DSB. Statistics were performed by MM. GM and BKP revised the manuscript and provided relevant intellectual content. All authors revised and approved the final draft.

---

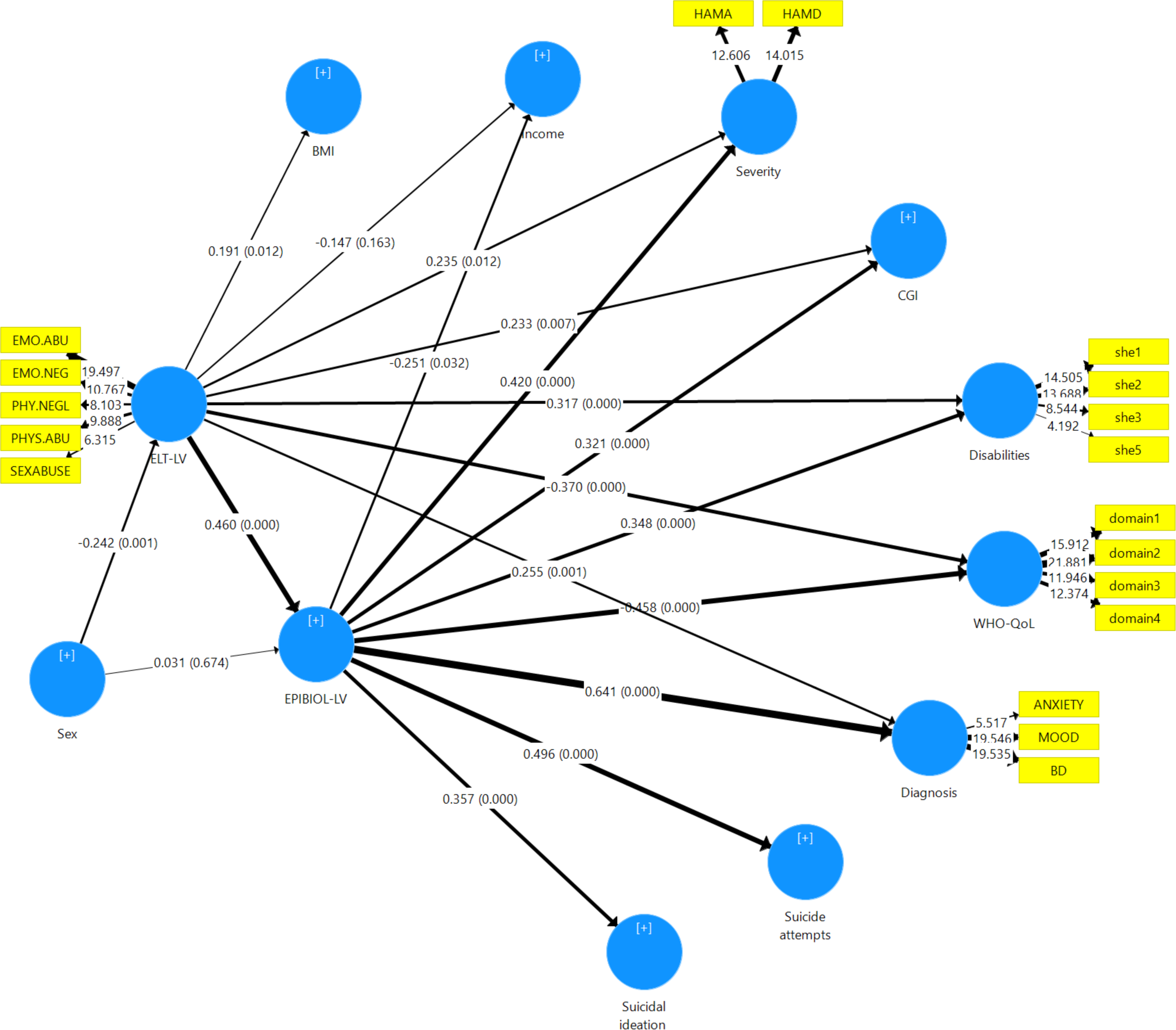

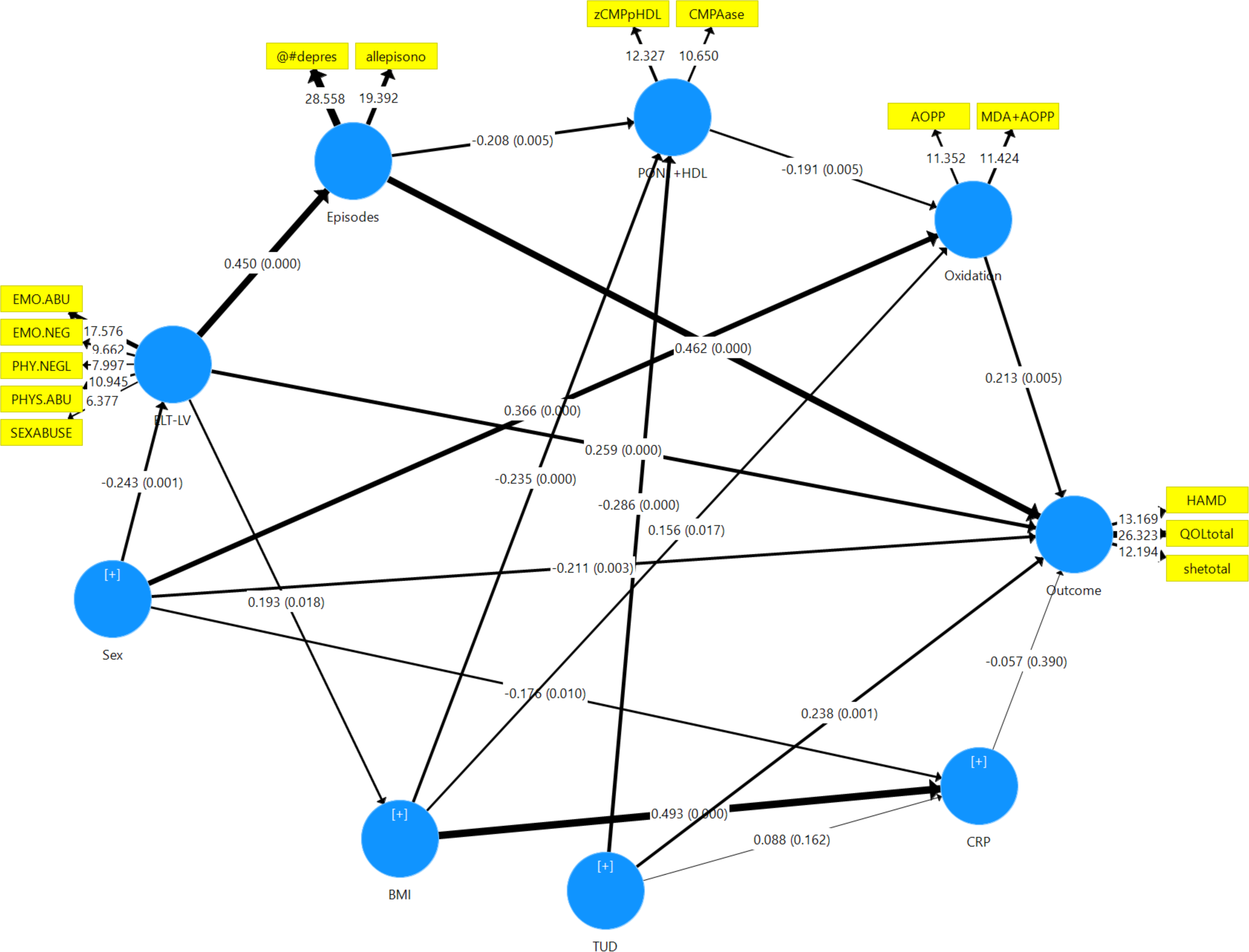

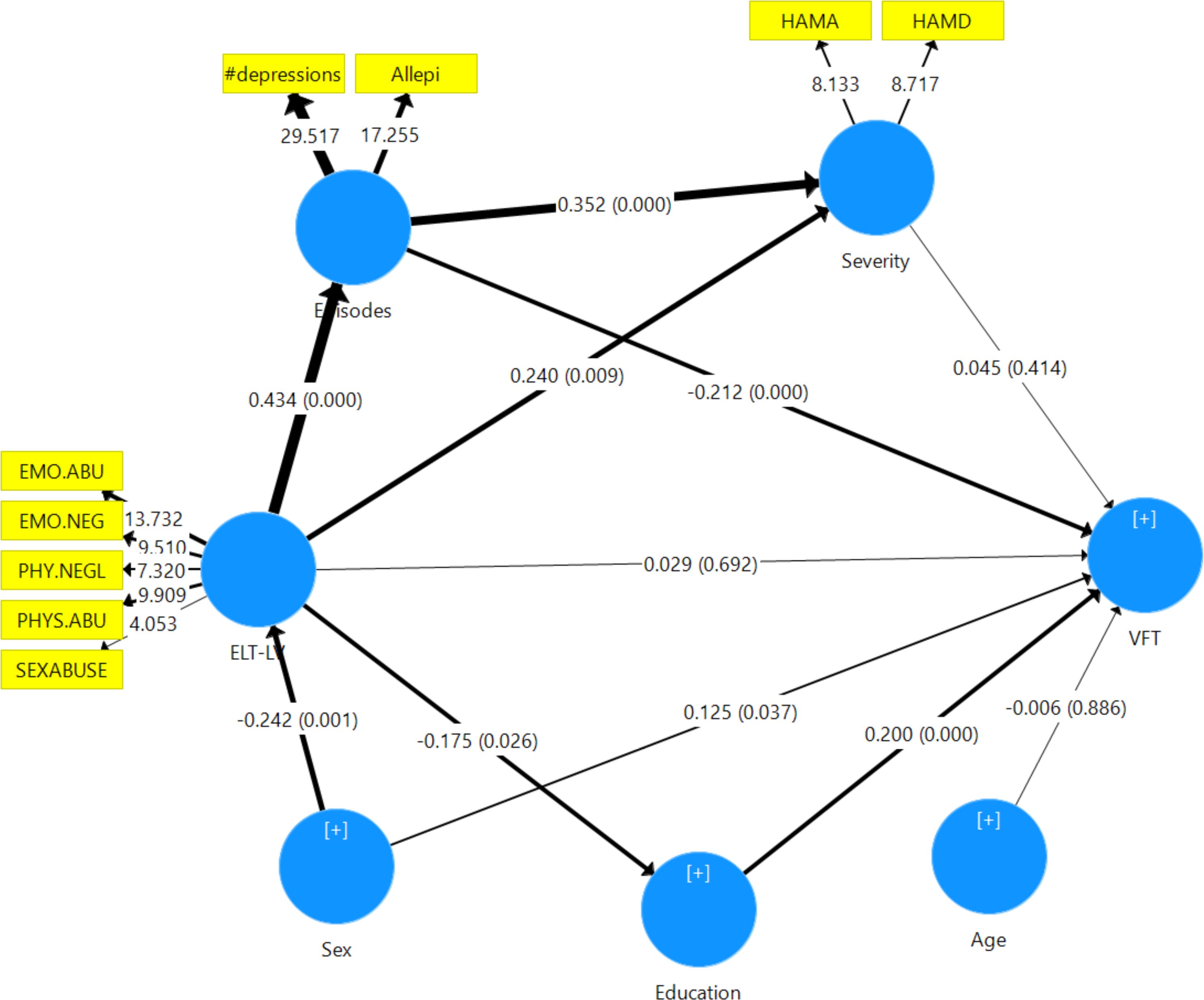

